# Comparative Analysis of MS/MS Search Algorithms in Label-Free Shotgun Proteomics for Monitoring Host-Cell Proteins Using Trapped Ion Mobility and ddaPASEF

**DOI:** 10.1101/2024.11.03.621185

**Authors:** Somar Khalil, Michel Plisnier

## Abstract

Host cell proteins (HCPs) are critical quality attributes that can impact the safety, efficacy, and quality of biotherapeutics. Label-free shotgun proteomics is a vital approach for HCP monitoring, yet the choice of tandem mass spectrometry (MS/MS) search algorithms directly influences identification depth and quantification reliability. In this study, six prominent MS/MS search tools—Mascot, MaxQuant, SpectroMine, FragPipe, Byos, and PEAKS—were systematically benchmarked for their performance on complex samples spiked with isotopically labeled proteins from Chinese hamster ovary cells, using trapped ion mobility spectrometry and parallel accumulation-serial fragmentation in data-dependent acquisition mode. Key performance metrics, including peptide and protein identifications, data extraction precision, fold-change (FC) accuracy, linearity, and measurement trueness, were evaluated. A Bayesian modeling framework with Hamiltonian Monte Carlo sampling was employed to robustly estimate FC means and variances, alongside local false discovery rates through posterior probability calibration. Bayesian decision theory, implemented via expected utility maximization, was used to balance accuracy against posterior uncertainty, providing a probabilistic assessment of each tool’s performance. Through this cumulative analysis, variability across tools was observed: some excelled in identification sensitivity and protein coverage, others in quantitative accuracy with minimal bias, and a few offered balanced performance across metrics. This study establishes a rigorous, data-driven framework for tool benchmarking, delivering insights for selecting MS/MS tools suited to HCP monitoring in biopharmaceutical development.

**Graphical abstract:** 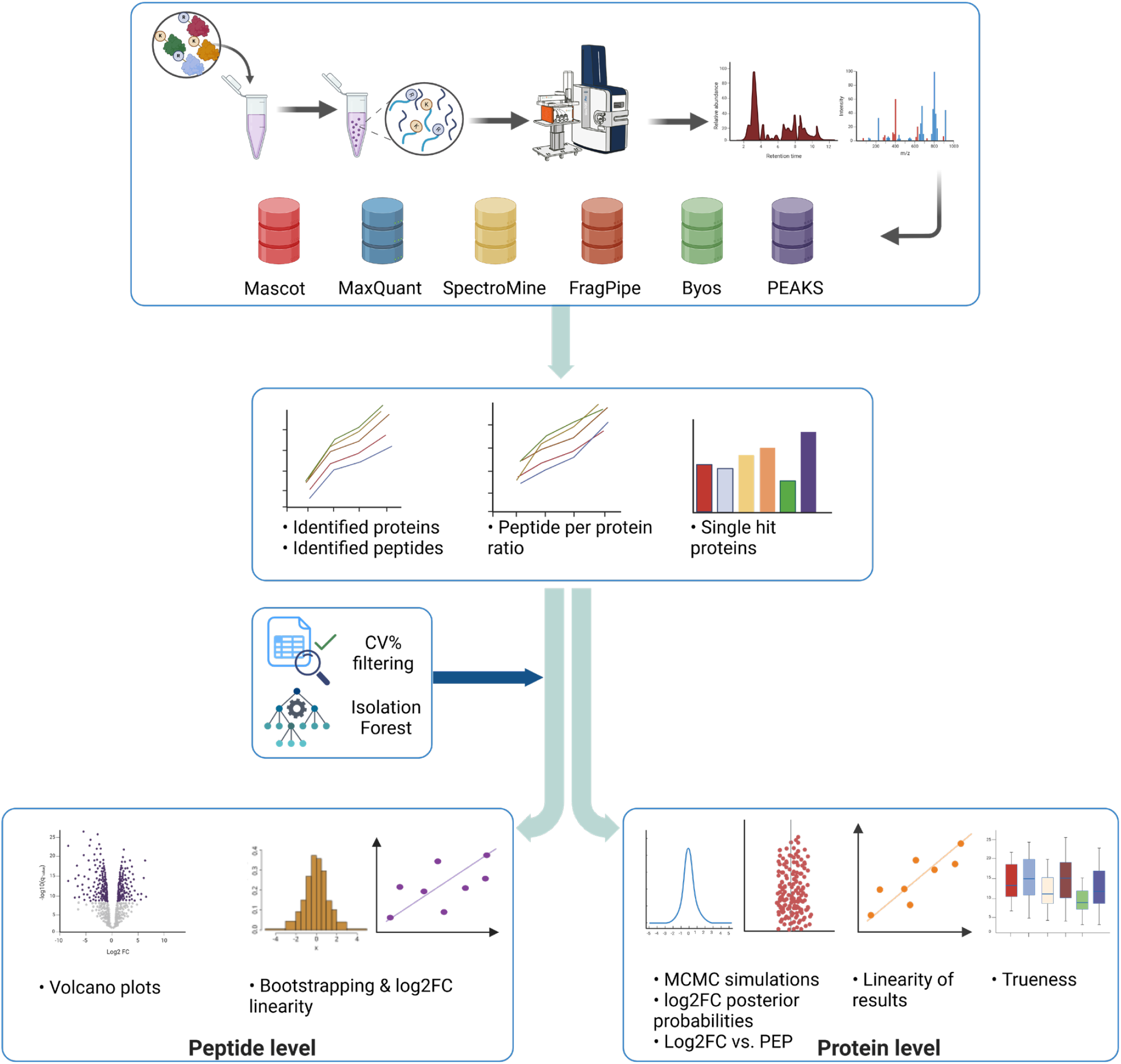

---

Host cell proteins (HCPs) are endogenous impurities that co-purify with target proteins during the manufacturing of recombinant therapeutics. The presence of HCPs poses risks to the safety and efficacy of biopharmaceutical products, necessitating rigorous monitoring and precise quantification^1-3^. Label-free quantification (LFQ) in shotgun proteomics has become a powerful tool for identifying and quantifying HCPs, yielding insights into their abundance and potential impacts on product quality^4-7^. However, the low abundance of HCPs and the broad dynamic range present significant detection challenges, primarily due to the stochastic nature of data-dependent acquisition (DDA) methods, which may affect the reliability and reproducibility of LFQ analyses. Recent advancements in tandem mass spectrometry (MS/MS) technologies, including parallel accumulation–serial fragmentation (PASEF) and trapped ion mobility spectrometry (TIMS), have demonstrated potential in addressing these limitations. PASEF enhances protein coverage and detection sensitivity by enabling the rapid collection of hundreds of fragmentation events per second^8-9^, while TIMS improves gas-phase resolution and selectivity by separating ions based on size, shape, and charge prior to MS/MS analysis^8-10^. These advancements can significantly improve the detection of low-abundance HCPs in complex biopharmaceutical samples. Beyond acquisition methods, the efficacy of LFQ shotgun proteomics largely depends on the MS/MS search tools employed for peptide sequence assignment, protein inference, and data extraction. The performance of these tools is closely tied to the underlying algorithms, scoring functions, and the implementation of non-parametric target-decoy competition (TDC) for false discovery rate (FDR) estimation at both peptide and protein levels^11-12^.

Prominent database search tools such as Mascot, MaxQuant, SpectroMine, and FragPipe are widely used in processing shotgun MS/MS data by matching experimental spectra against theoretical spectra generated from *in-silico* protein digestions^13^. These tools utilize diverse scoring functions, which typically assign probabilities to peptide-spectrum matches (PSMs) and convert them into scores through generative models based on probability distributions for spectrum-peptide classification^14-15^. For instance, Mascot employs a probability-based scoring model derived from the MOWSE algorithm to rank PSMs according to peptide molecular weight distributions and integrates a hypergeometric model for protein-level FDR estimation, similar to the MAYU approach^16-19^. MaxQuant, in contrast, utilizes the Andromeda algorithm, which models fragment ion match probabilities with a binomial distribution^20^. Posterior error probabilities (PEP) for PSMs are subsequently derived by comparing scores against decoy distributions, yielding confidence measures for each match^21-22^. SpectroMine utilizes the Pulsar search engine, which integrates the probabilistic mProphet model for scoring, enhanced by support vector machine (SVM) classification for PSM rescoring^23^. FragPipe, employing the MSFragger algorithm, accelerates PSM identifications through fragment ion indexing and refines scoring with mass accuracy considerations^24-25^. For PSM rescoring and validation, FragPipe uses PeptideProphet (a mixture model-based PEP estimator) or Percolator (an SVM-based classifier)^26-27^.

Proteins are then inferred by ProteinProphet, which groups peptides and assigns probabilities at the protein level^28^. Hybrid approaches that integrate elements of *de novo* sequencing with database searches are exemplified by tools such as PEAKS and Byos. PEAKS extracts sequence tags to shortlist proteins from the database, followed by an error-tolerant search to match flanking masses^29-32^. It employs a dot-product similarity search for the scoring function and a modified TDC approach, termed “decoy fusion”, in which target and decoy sequences are concatenated prior to protein inference^33^. Byos, incorporating the Byonic algorithm, leverages flanking masses (lookup peaks) to extract candidate peptides^34-35^. The scoring function also utilizes a dot-product algorithm, and spectral identifications are further integrated through the ComByne probabilistic protein assembly tool^35-36^. Furthermore, Byos implements a “ProteinFirst” strategy, applying a two-dimensional TDC to control FDR at both PSM and protein levels simultaneously^37^.

In this complex analytical landscape, Bayesian inference can be a powerful approach for quantifying uncertainties in protein identification and differential abundance analyses^38^. Among various Markov Chain Monte Carlo methods, Hamiltonian Monte Carlo (HMC) is particularly effective in navigating high-dimensional parameter spaces by leveraging gradient information from posterior distributions^39-40^, thereby reducing random walks, and increasing the precision of posterior variance estimates in protein quantification. Additionally, Bayesian PEP values can offer probabilistic confidence measures for the differentially abundant proteins^38,41^.

This study comprehensively benchmarks six widely used MS/MS search tools—Mascot, MaxQuant, SpectroMine, FragPipe, Byos, and PEAKS—to evaluate their performance in identifying and quantifying HCPs within complex biopharmaceutical samples. Antigen protein samples spiked with stable-isotope-labeled proteins from Chinese hamster ovary (CHO) cells were analyzed using TIMS and ddaPASEF. Key performance metrics, including peptide and protein identifications, data extraction precision, fold-change (FC) accuracy, linearity, and trueness were rigorously assessed. Bayesian inference, leveraging HMC sampling and Bayesian FDR control, was central to the posterior estimation of log2FC errors and confidence at the protein level. Bayesian decision theory (BDT), implemented through an expected utility maximization framework, was applied to balance FC accuracy against posterior uncertainty. By systematically evaluating the capabilities and limitations of these tools, this study delivers a comprehensive, data-driven assessment of their reliability in monitoring HCPs within an LFQ framework, tailored to the biopharmaceutical industry.

## EXPERIMENTAL SECTION

### Materials

A recombinant antigen drug substance (DS) was provided in-house from GSK (Rixensart, Belgium). Trypsin/Lys-C protease mix, peptide desalting spin columns, dithiothreitol (DTT), triethylammonium bicarbonate (TEAB), and iodoacetamide (IAM) were purchased from Thermo Scientific, Waltham, MA, USA. MassPREP Bovine Serum Albumin (BSA, SwissProt P02769) digestion standard, MassPREP Alcohol Dehydrogenase (ADH, SwissProt P00330) digestion standard, MassPREP Phosphorylase b (PYGM, SwissProt P00489) digestion standard, MassPREP Enolase (ENO1, SwissProt P00924) digestion standard, and RapiGest SF surfactant were purchased from Waters, Milford, MA, USA. SILu-CHOP stable-isotope labeled CHO proteins (MSQC12) was purchased from Sigma-Aldrich, Overijse, Belgium. Formic acid (FA), trifluoroacetic acid (TFA), acetonitrile (ACN), and LC-MS grade water from Biosolve, Valkenswaard, Netherlands. Evotip trap columns and EV-1137 Performance C18 column (1.5 μm, 150 μm x 150 mm) were purchased from Evosep, Denmark.

### Proteomics sample preparation and digestion

DS samples, each containing 20 μg, were spiked with the SILu-CHOP standard at four levels: 1 μg, 2 μg, 3 μg, and 4 μg (referred to as L1, L2, L3, and L4, respectively). Each spiked sample was prepared in triplicate. Samples were denatured and reduced using 0.1% RapiGest and 8 mM DTT for 45 min at 50 °C. Alkylation was carried out with 16 mM IAM for 30 min in the dark at room temperature. Enzymatic digestion was then performed overnight at 37 °C using a trypsin/Lys-C mix at an enzyme-to-protein ratio of 1:40. Following digestion, the samples were acidified with 0.5% TFA and centrifuged to collect the supernatants. These were subsequently desalted using spin columns and dried using a speed vacuum concentrator. The dried samples were reconstituted in 0.1% FA and spiked with the four MassPREP protein digest standards (0.30 fmol/μL, 0.80 fmol/μL, 1.24 fmol/μL, and 4.0 fmol/μL for PYGM, ADH, ENO1, and BSA, respectively). Each sample (500 ng) was loaded into Evotip trap columns according to the manufacturer’s instructions.

### Liquid Chromatography-Mass Spectrometry Analysis

Samples loaded onto Evotip trap columns were separated in a 44-min gradient (30 samples per day) on an EV-1137 Performance column with the Evosep One system (EV-1000, Evosep, Denmark). The mobile phase A consisted of 0.1% FA in water, while mobile phase B was 0.1% FA in ACN. Peptide separation was conducted at 40°C with 35% solvent B at a flow rate of 500 nL/min. The Evosep One system was coupled to a timsTOF Pro MS (Bruker Daltonics, Germany) via a CaptiveSpray nano-electrospray ion source. ddaPASEF mode was utilized, with eight PASEF ramps and a mass-to-charge (m/z) range of 100–1350. Singly charged precursor ions were systematically excluded using the polygon filter. The acquisition settings included a target intensity of 24,000 and an intensity threshold of 1,200. A 1.0-second cycle time was applied, with TIMS configured for 100 ms for both ramp and accumulation times. Collision energy was ramped linearly as a function of mobility, with default isolation widths applied. Active exclusion was maintained with a 0.5-min release time. TIMS calibration was performed linearly to determine reduced ion mobility (IM) coefficients (1/K_0_), using calibration ions with m/z and 1/K_0_ values of 622.02 at 0.98 V/cm^2^, 922.01 at 1.19 V/cm^2^, and 1221.99 at 1.38 V/cm^2^. IM range was set from 0.7 to 1.4 V/cm^2^.

### Data processing

All ddaPASEF raw data were processed against an in-house protein database containing 5280 protein sequences of *C*.*griseus* entries, sourced from UniProt (Swiss-Prot and TrEMBL annotations). The global search parameters were set to include trypsin/P specificity, allowing for one missed cleavage, with precursor mass tolerance of 15 ppm and fragment mass tolerance of 30 ppm. The fixed modifications included carbamidomethylation of cysteine, [^13^C_6_, ^15^N_4_]-Arginine and [^13^C_6_, ^15^N_2_]-Lysine, while methionine oxidation was considered a variable modification. A separate search was performed for the four standard protein digests of MassPREP (P02769, P00330, P00489, and P00924), with carbamidomethylation as the sole fixed modification. Raw data were searched with Mascot (v2.7) using the decoy option, and Mascot identification results containing PSMs were imported into Skyline (v21.1) for peak integration and peptide quantification using extracted ion chromatograms (XICs) from MS1 precursor ions. For simplicity, the combined usage of Mascot and Skyline for the quantification is referred to as Mascot in the study. MaxQuant (v2.6) was used with its default TIMS-DDA settings and included four-dimensional isotope pattern matching between runs (MBR) with windows set at 0.7 min for retention time and 0.05 1/K_0_ for IM. A reverted decoy database was employed, and an FDR of 1% was maintained at both the PSM and protein levels. With SpectroMine (v4.5), a q-value filter of 0.01 at the precursor, peptide, and protein levels was applied, with default settings otherwise. FragPipe (v22.0) workflow used the MSFragger (v4.1) engine with default settings, integrating Percolator for PSM validation (minimum probability of 0.5) and ProteinProphet for protein inference, also applying a 1% FDR threshold. IonQuant^42^ MBR feature was enabled with 0.7 min and 0.05 1/K_0_ tolerances. PEAKS Studio (v12.0) processed the raw data using the PEAKS DB workflow, with parameters optimized for the TimsTOF instrument using CID fragmentation. The protein FDR was set to 1%, and peptide scores were required to be at least 20 (-10logP). MBR parameters included a 15-ppm mass error tolerance, automatic retention time shift tolerance, and a 0.05 1/K_0_ IM tolerance, with decoy sequences generated via the PEAKS decoy-fusion method. Finally, Byos (v5.7) was employed, setting a protein cut-off at 1% FDR and a |Log Prob| score at least 2.0 lower than the top decoy protein score. PSM filtering was performed post-protein assembly, using a 15-ppm extraction window and a 1.5-min XIC search window. For quantification, only peptides below 0.01 for protein-wise two-dimensional PEP and two-dimensional FDR were accepted.

## RESULTS AND DISCUSSION

Six commonly used MS/MS search algorithms were compared to evaluate their performance in processing MS/MS data generated by spiking stable-isotope-labeled CHO proteins into a purified antigen DS at four spike levels. Each benchmark sample was prepared in technical triplicates and analyzed in ddaPASEF mode. Statistical analyses, Bayesian inference modeling, and data visualization were conducted using Python (v3.6), R (v4.3), and GraphPad Prism (v10.3; GraphPad Software, San Diego, USA).

### Performance of labeled CHO proteome identification

To evaluate the efficacy of each MS/MS search tool, common contaminants and peptides shorter than seven amino acids were excluded from protein and peptide identifications. As shown in **Figure 1A**, the number of identified CHO proteins increased proportionally with the spike level across all tools. PEAKS exhibited the highest protein coverage, with identifications rising from 472 proteins at L1 to 1565 proteins at L4, totaling 1634 unique proteins. SpectroMine and FragPipe also demonstrated strong protein identification capabilities, with cumulative identifications of 1434 and 1466 proteins, respectively. In contrast, Byos, MaxQuant and Mascot displayed lower identification counts, identifying between 1054 and 1248 proteins. **Figure 1B** illustrates peptide-level identifications. Byos outperformed all other tools with a total of 9486 peptides, followed closely by SpectroMine, PEAKS, and FragPipe, which identified between 7851 and 8342 peptides. MaxQuant and Mascot identified fewer peptides, with totals of 4971 and 5118, respectively. Peptide-to-protein ratios, depicted in **Figure 1C**, indicate that Byos achieved the highest ratios, ranging from 4.8 to 6.7 across spike levels, suggesting greater depth for identified proteins. In contrast, MaxQuant and Mascot exhibited the lowest ratios. Single-hit proteins, shown in **Figure 1D**, varied significantly among the tools. PEAKS reported the highest number of single-hit proteins (569), followed by FragPipe (483) and SpectroMine (364). Byos, however, had the fewest single-hit proteins at 181. The UpSet plots (**Figure 1E, Figure 1F**) illustrate the overlap in identified proteins and peptides across the tools. Notable intersections exist, highlighting shared identifications among tools despite distinct algorithmic approaches related to the heuristic protein inference. PEAKS identified 102 unique proteins, while Byos reported 933 unique peptides. Overall, SpectroMine, FragPipe, PEAKS, and Byos demonstrated superior sensitivity across both protein and peptide coverage, effectively capturing low-abundance CHO proteins at all spike levels. Conversely, Mascot and MaxQuant exhibited comparatively lower identification counts and peptide-to-protein ratios, particularly at lower spike levels.

**Figure 1.**
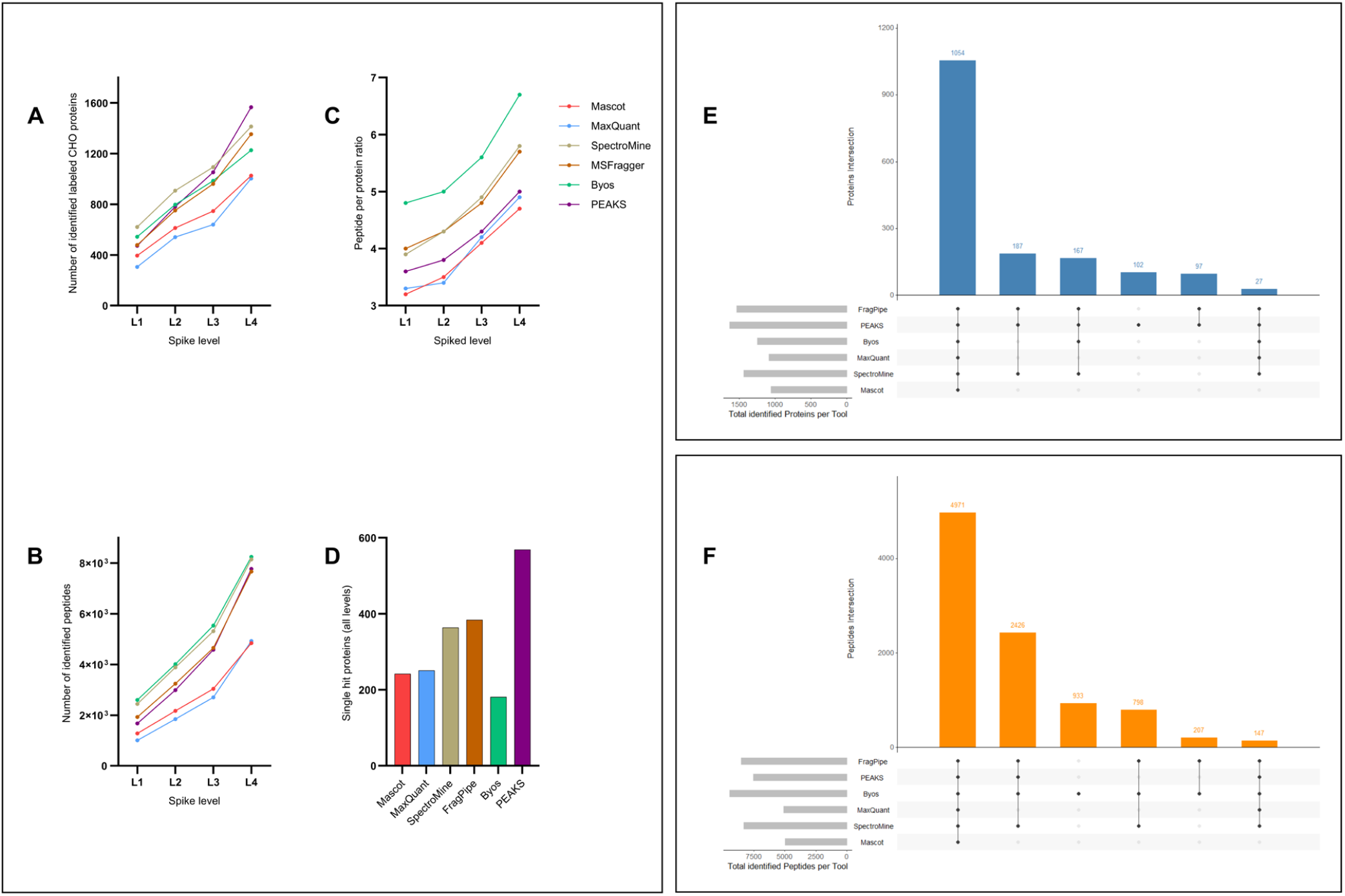
Performance comparison of the six MS/MS search tools in identifying labeled CHO proteins and peptides across four spike levels (L1-L4). Number of identified labeled CHO proteins (**A**) and peptides (**B**) at each spike level. (**C**) Peptide-to-protein ratios. (**D**) Number of single-hit proteins. UpSet plot showing the intersection of identified labeled CHO proteins (**E**) and peptides (**F**) by unique or shared combinations across tools.

### Performance of labeled CHO proteome quantification

Prior to downstream analysis, several data filters were applied. Only doubly and triply charged peptides were retained, while single-hit proteins and modified peptides were excluded. For each spike level, peptides without missing values across triplicates were selected. To normalize XIC intensities, each peptide’s intensity was divided by the total intensity of its respective replicate, then scaled by the mean total intensity across triplicates. Datasets were log10-transformed and subjected to outlier detection using the Isolation Forest algorithm^43^, an unsupervised decision tree-based method, with an anomaly threshold of 0.55. Peptides exhibiting irregular intensity patterns were identified as outliers and excluded from further analyses (**Supporting Information, Figure S1**). To evaluate data extraction precision, the coefficient of variation (CV%) was calculated for each tool across the four spike levels. The log10-transformed CV% of each peptide was plotted against its log10-transformed geometric mean intensity, as shown in **Figure 2A**. FragPipe consistently demonstrated superior data extraction precision, with the majority of peptide CV% values falling below the 25% threshold, indicated by the red horizontal line. This performance was further reflected in the lowest median CV% values across spike levels (10.9–12.6%). SpectroMine exhibited similar performance, with median CV% values ranging from 13.6% to 14.5%, underscoring its reproducible data extraction. Byos (14.4-18.7%), PEAKS (15.3-18.1%), and Mascot (16.6-19.5%) displayed moderate precision. In contrast, MaxQuant showed the highest degree of variability, particularly at lower spike levels, with CV% values spanning from 23.4% to 34.8% (**Supplementary Information, Table S1**).

**Figure 2.**
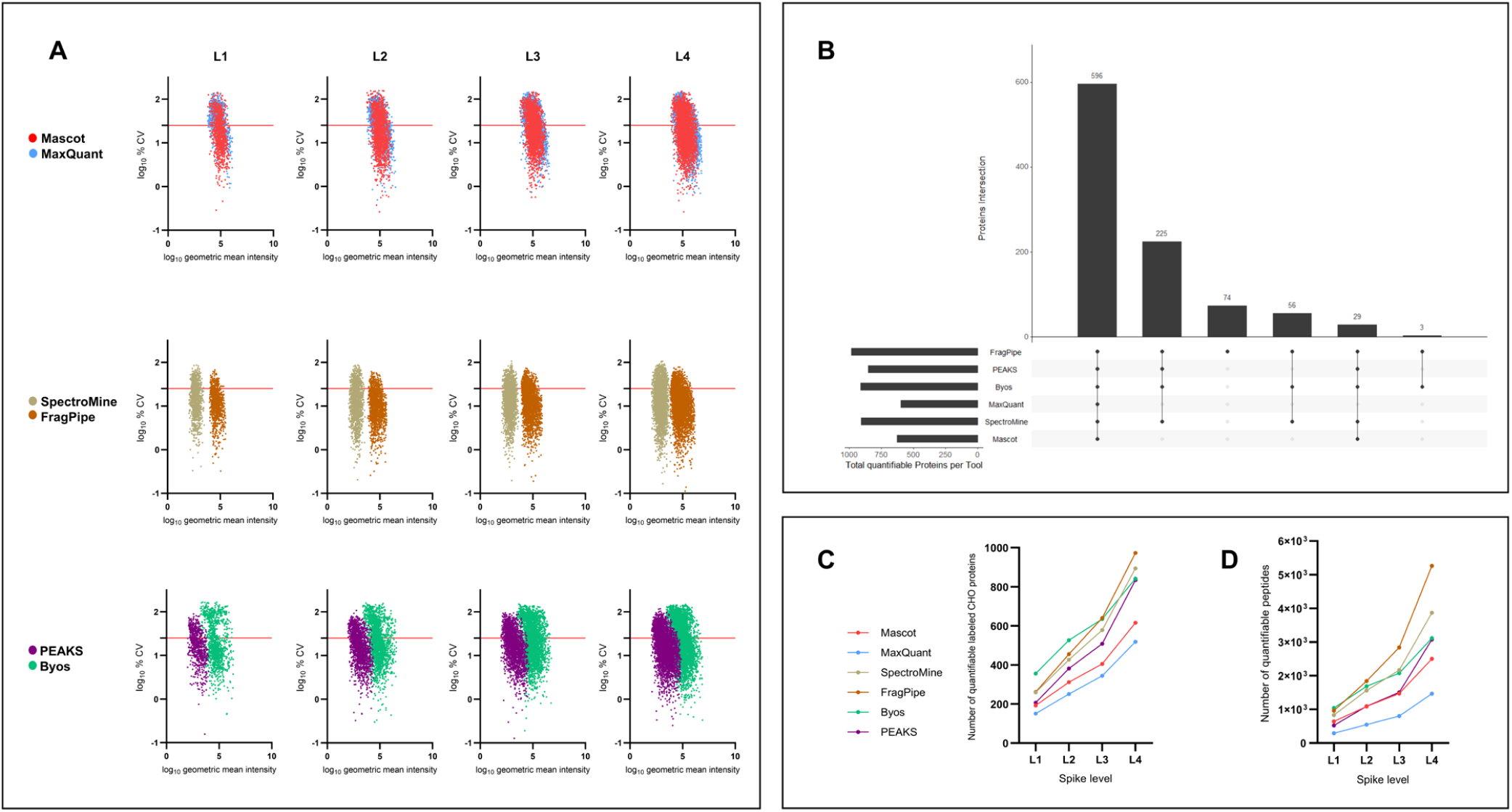
Performance comparison of the six MS/MS tools for data extraction precision and quantifiable labeled CHO proteins and peptides across four spike levels. (**A**) Scatter plot of CV% vs. geometric mean intensity (both log10-transformed) for each spike level and MS/MS tool. Each point represents a peptide, with color coding for each MS/MS tool. Horizontal red lines indicate the threshold of 25% CV (log10-transformed). (**B**) UpSet plot illustrating the intersection of quantifiable labeled CHO proteins by unique or shared combinations of MS/MS tools. Number of quantifiable labeled CHO proteins (**C**) and peptides (**D**) at each spike level.

#### Peptide level

Peptide intensities with a CV% exceeding 25% across replicates were excluded, and the remaining peptides were classified as quantifiable peptides (**Figure 2D**). FragPipe, which demonstrated consistently low CV%, reported the highest number of quantifiable peptides, ranging from 962 at the lowest to 5265 at the highest spike level. Byos (1043-3117) and SpectroMine (832-3869) followed closely, with Byos showing a slight advantage at lower spike levels. Mascot (640–2501) and PEAKS (520–3076) exhibited moderate performance, both improving as spike levels increased. MaxQuant quantified the fewest peptides, with counts ranging from 290 to 1465. Overall, FragPipe, Byos, and SpectroMine outperformed the other tools, particularly at lower spike levels.

To evaluate the statistical significance and FC accuracy of the quantifiable peptides, an unpaired Student’s t-test was conducted with the adaptive linear step-up method of Benjamini, Krieger, and Yekuteli^44^ for multiple hypothesis correction. Six pairwise comparisons were performed: L2 vs. L1, L3 vs. L1, L3 vs. L2, L4 vs. L1, L4 vs. L2, and L4 vs. L3, with expected log2FC values of 1.00, 1.58, 0.58, 2.00, 1.00, and 0.41, respectively. Due to the high level of missingness at lower spike levels, missing peptide intensity values were not imputed to prevent potential bias. Volcano plots were generated (**Figure 3A**), with log2FC values deemed statistically significant at a q-value threshold of <0.05. Additionally, median absolute log2FC errors between expected and observed values were calculated for each comparison level (**Supporting Information, Table S2**). Quantitative performance at the peptide level was assessed by analyzing the distribution of negative log10-transformed q-values, the density of peptides aligning with the expected log2FC trendlines, and the median absolute log2FC errors. In the L2 vs. L1 and L3 vs. L1 comparisons, Mascot, MaxQuant, FragPipe, and Byos showed comparable performance, with a high density of peptides clustering around the expected log2FC values. Byos displayed a slightly wider dispersion of data points relative to the expected trendline, while Mascot reported the lowest median absolute log2FC errors (0.20–0.23). In the L3 vs. L2 comparison, all tools demonstrated good performance, though PEAKS and FragPipe showed slight underperformance. In comparisons involving L4, SpectroMine exhibited superior differential accuracy, evidenced by low median absolute log2FC errors (0.25–0.49). Byos followed closely, showing reasonable alignment and balanced dispersion around the expected trendline. Conversely, FragPipe and PEAKS demonstrated the poorest performance, with skewed dispersion toward positive deviations from the expected trendline and the highest median absolute log2FC errors. This pattern suggests an overestimation of peptide intensities relative to their expected differential abundance.

**Figure 3.**
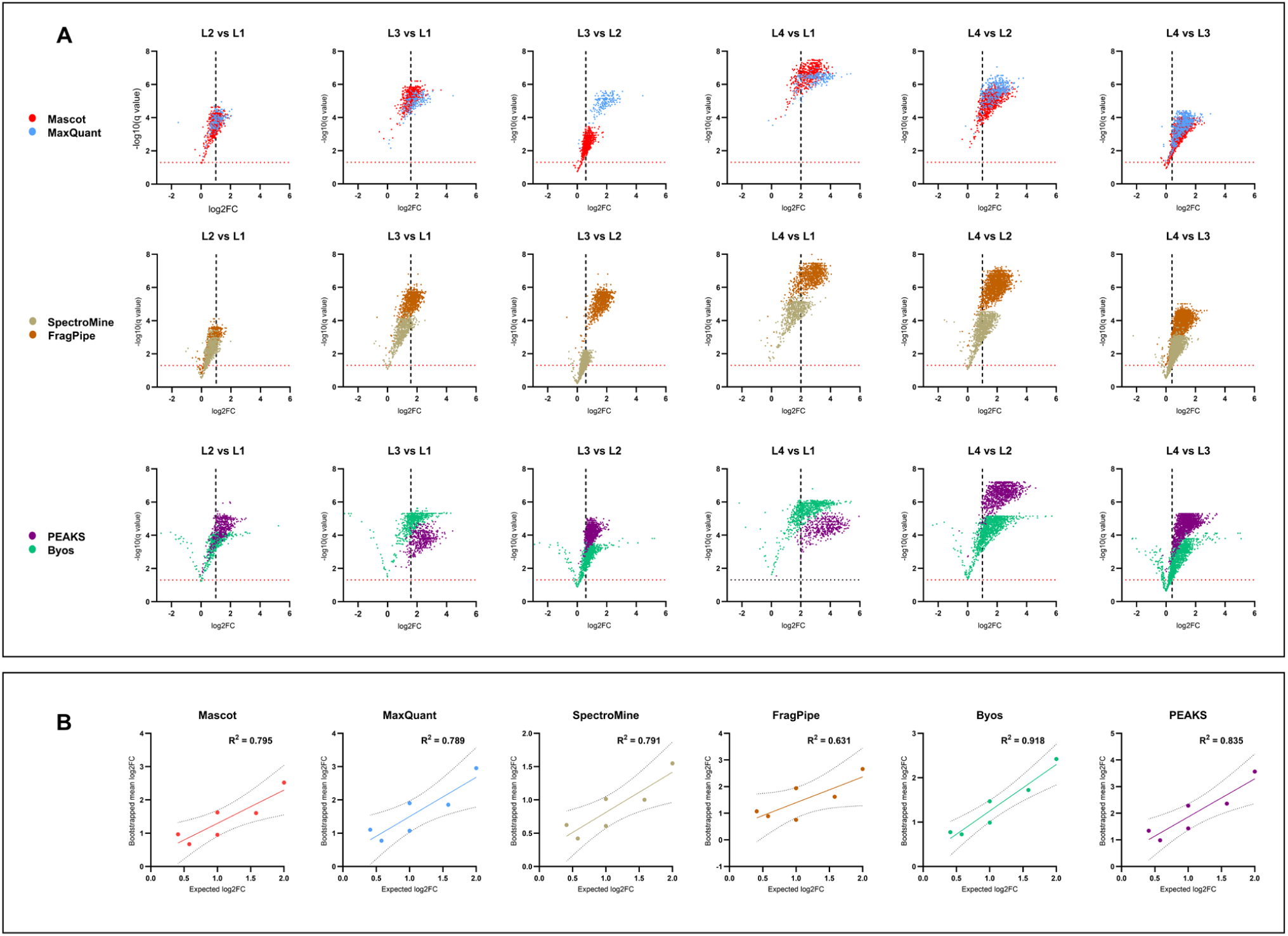
Evaluation of log2FC accuracy at the peptide level across six MS/MS tools in comparison to expected values. (**A**) Volcano plots for each tool showing the significance (-log10 q-value) versus log2FC for six spike level comparisons (L2 vs L1, L3 vs L1, L3 vs L2, L4 vs L1, L4 vs L2, and L4 vs L3). The vertical dashed lines represent expected log2FC values, and the horizontal red line indicates the significance threshold (q-value = 0.05). Each point represents a peptide, with more points aligned along the expected log2FC indicating better accuracy. (**B**) FC regression plots showing the correlation between expected and bootstrapped mean log2FC values for each MS/MS tool across spike level comparisons. R^2^ values reflect the linearity of the relationship.

Furthermore, a bootstrapping method was employed to estimate the mean log2FC for each comparison level across all tools, utilizing resampling with replacement. Through 6000 bootstrap iterations, log2FC values for each comparison level were repeatedly resampled, and the mean log2FC was calculated for each iteration. The overall mean of these bootstrap means was then computed, providing a robust, distribution-free estimate of mean log2FC while minimizing the influence of outliers and random fluctuations. Subsequently, a linear regression model was fitted for each tool to assess the correlation between the bootstrapped means and expected log2FC values. The regression lines and corresponding coefficients of determination (R^2^), visualized in **Figure 3B**, offer a comparative analysis of each tool’s ability to capture expected FCs linearly. Byos exhibited the highest R^2^ value of 0.918, indicating a strong linear correlation with minimal impact from outliers. Interestingly, PEAKS followed closely, achieving an R^2^ of 0.835. This moderate performance, despite displaying the highest log2FC errors, indicates that PEAKS maintained a proportional FC bias across the full range of comparisons. Mascot, SpectroMine, and MaxQuant yielded comparable R^2^ values of 0.795, 0.791, and 0.789, respectively, suggesting reasonable linearity with modest data dispersion around the regression lines. In contrast, FragPipe displayed the lowest R^2^ value at 0.631, reflecting a weaker correlation likely due to disproportionate FC errors across the comparisons and highlighting challenges in maintaining linear differential estimation at the peptide level.

#### Protein level

The UpSet plot in **Figure 2B** illustrate a significant overlap of quantifiable proteins across the tools, with FragPipe reporting 74 unique proteins. As depicted in **Figure 2C**, Byos exhibited robust performance, with quantifiable protein counts increasing from 356 at the lowest spike level (L1) to 841 at the highest spike level (L4). SpectroMine and FragPipe also demonstrated strong capabilities, particularly at higher spike levels, with FragPipe reaching a peak of 973 quantifiable proteins at L4. In contrast, MaxQuant and Mascot reported comparatively lower quantification, with 518 and 616 proteins quantified at L4, respectively. PEAKS, which had the lowest initial count at L1 (207 proteins), increased to 834 quantifiable proteins at L4. This trend suggests variability in tool efficiency, with FragPipe, Byos, and SpectroMine achieving the highest numbers of quantifiable proteins.

The Hi3 LFQ approach was applied across the four spike levels to derive quantifiable protein intensities. Protein intensities were calculated from the top three most intense quantifiable peptides at each spike level using sum, average, and median as aggregation methods. A Bayesian framework employing HMC sampling was implemented to model the log2FC between spike levels. A Student’s t-distribution likelihood function with three degrees of freedom was chosen to account for potential outliers. Weakly informative priors were placed on the mean log2FC (*μ*), centered around the expected log2FC values, while half-normal priors were applied to the scale parameter (*σ*), capturing uncertainty around (*μ*). The model was fitted independently for each comparison using the three aggregation methods. The HMC sampling was executed with the No-U-Turn Sampler, an adaptive algorithm that optimizes sampling by dynamically adjusting parameters for efficient exploration of the posterior distribution^45^. Six chains with 8000 iterations each were employed to ensure convergence, with model convergence assessed via the *R-hat* statistic^46^. Posterior distributions of log2FCs were derived for each tool across comparisons, providing a probabilistic interpretation based on observed data and prior expectations. Additionally, PEP values were computed, and the correlation between log2FC and PEP was plotted to quantify the uncertainty and assess the statistical reliability of the FC estimates for each tool. Furthermore, to systematically balance accuracy and uncertainty, BDT was applied within an expected utility maximization framework. A utility function (*U*) was defined as shown in Eq. 1:

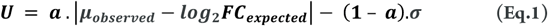

Where *a* = 0.7 was selected to prioritize accuracy over uncertainty. This expected utility function ranks the MS/MS tools by maximizing the expected posterior utility, integrating both differential accuracy and precision into a comprehensive assessment of tool performance.

Significant differences in posterior results were observed depending on the protein aggregation method. For SpectroMine, the sum aggregation demonstrated better alignment with the expected log2FC values, as evidenced by narrower posterior distributions and lower values of (*σ*). Conversely, the remaining five tools yielded more accurate results with the average or median aggregation methods (**Supporting Information, Figure S2**). To ensure fairness in the benchmark, the sum aggregation was applied for SpectroMine, while the average aggregation was used for the other MS/MS tools.

Posterior probability distributions, visualized as density plots in **Figure 4A**, illustrate the mean, uncertainty, and the 95% posterior credible intervals for each tool. While narrow credible intervals were observed across most comparison levels, log2FC accuracy varied among tools. Byos, SpectroMine, and Mascot displayed superior performance, with posterior distributions tightly centered around the expected log2FC values and overall median errors of 0.12, 0.14, and 0.21, respectively (**Supporting Information, Table S3**). MaxQuant and FragPipe exhibited comparable performance, with median errors of 0.45 and 0.47, respectively, while PEAKS (0.70) showed the largest deviations from expected log2FC values.

**Figure 4.**
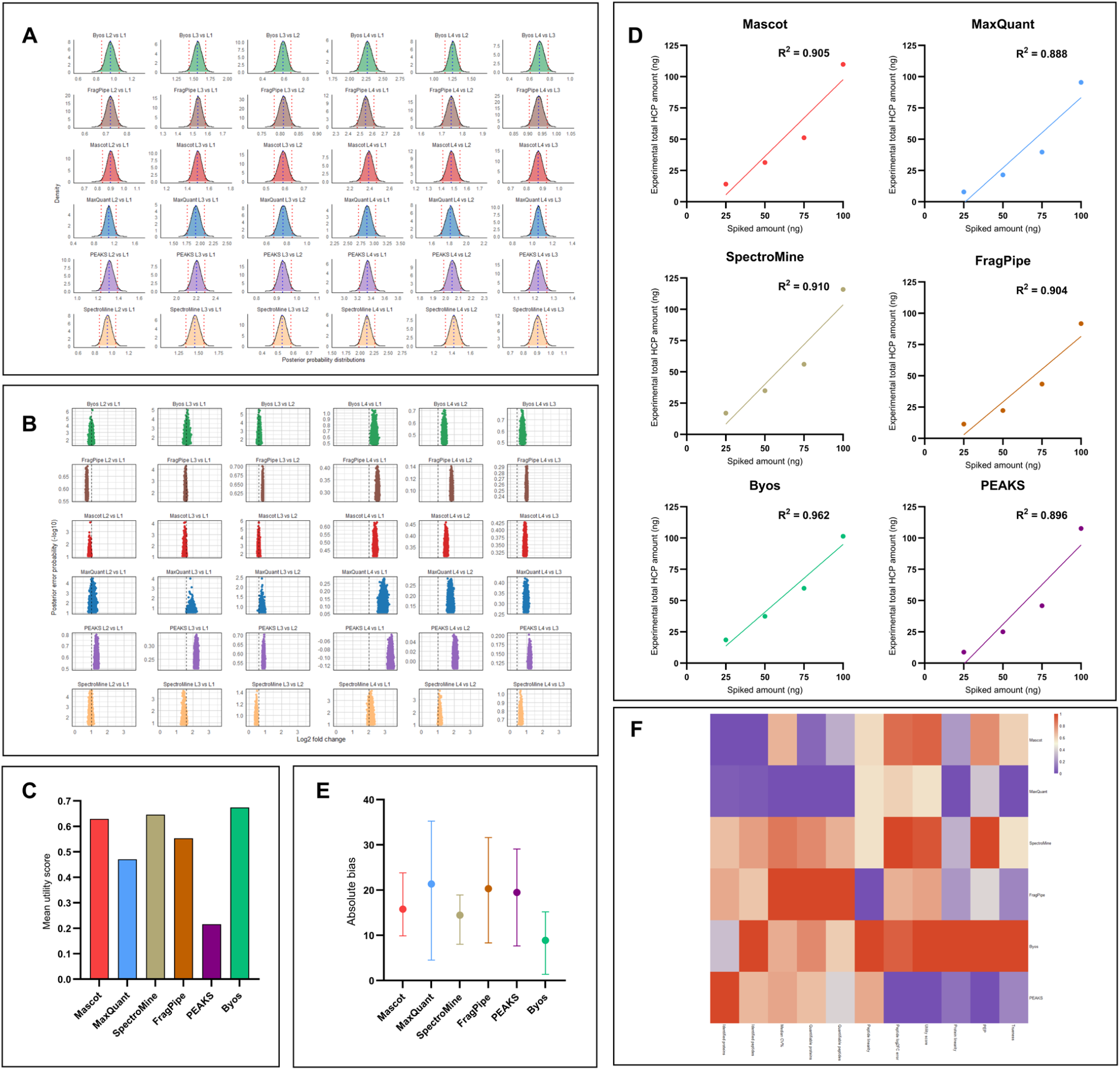
(**A**) Density plots for posterior probability distributions of log2FC values estimated by different tools across comparisons. Vertical blue dashed lines indicate the posterior means. Vertical red dotted lines represent the 95% credible intervals. (**B**) PEP vs. log2FC plots providing a statistical assessment of the log2FC estimates against PEP, illustrating the relationship between the measured log2FC and its associated uncertainty. Points distribution along the x-axis indicates the magnitude of change, while their position on the y-axis reflects the -log10(PEP) values. (**C**) Bar chart of mean utility scores derived from BDT for all MS/MS tools. Scores are calculated based on a combination of log2FC accuracy and variance with (*a*) value of 0.7 at each comparison level. (**D**) Linearity of observed vs. expected overall HCP abundance across MS/MS tools. Each tool’s performance is visualized with a line of best fit. The solid-colored line represents the observed trend for each tool. (**E**) Mean absolute bias with range across MS/MS tools represented by bars. The absolute bias is the absolute difference between the observed and expected overall HCP abundance values at different spike levels (25 ng, 50 ng, 75 ng, 100 ng). (**F**) Comparative heatmap illustrating the benchmarking of MS/MS tools across all performance metrics. Each row representing an individual tool and each column corresponding to a distinct, scaled performance metric. Evaluated metrics include protein and peptide identifications, median CV%, quantifiable proteins and peptides, bootstrapped differential peptide linearity (R^2^ values), peptide median log2FC errors, mean utility scores, protein linearity of results (R^2^ values), mean -log10(PEP) values, and trueness (absolute bias). All metrics were scaled to facilitate comparability, with a color gradient ranging from blue (lower performance) through white to red (higher performance).

Log2FC vs. PEP plots in **Figure 4B** further corroborated these trends. Byos and SpectroMine demonstrated strong alignment with expected values, producing more confident FC estimates, as indicated by lower PEP values and the highest mean -log10(PEP) scores across all comparisons (**Supporting Information, Table S4**). Mascot performed well in certain comparisons but exhibited variability. FragPipe, MaxQuant, and PEAKS showed higher PEP values, with PEAKS demonstrating the weakest performance overall. Utility function, calculated through BDT, reinforced these findings (**Figure 4C**). Byos, SpectroMine, and Mascot ranked highest, with mean *U* values of 0.67, 0.65, and 0.63, respectively. PEAKS, exhibiting larger deviations and higher posterior variance, attained a mean *U* value of 0.22, reflecting its limitations in producing reliable differential analysis, particularly for comparisons involving smaller expected FCs. Posterior predictive checks (**Supporting Information, Figure S3**) indicated a satisfactory fit between observed and simulated FCs, although slight discrepancies were observed in L4 comparisons for certain tools. Lastly, a sensitivity analysis on the (*a*) parameter in the utility function was performed, varying between 0.5 and 0.8. Results demonstrated consistent overall rankings, with Byos retaining the highest position and SpectroMine showing slight improvement at lower (*a*) values, attributable to its lower variance.

#### HCP Hi3 LFQ quantification

The top three peptide intensities of the four proteins from the MassPREP standards were aggregated using the sum for SpectroMine and the average for the other tools. These derived MassPREP protein intensities were plotted against their theoretical loads (fmol) to construct a universal signal-response curve^4, 47^(**Supporting Information, Figure S4**). This linear curve was used to estimate the individual abundance of labeled CHO proteins (in fmol), which were subsequently converted to ng values based on molecular weights. The performance of the MS/MS tools was then evaluated in terms of linearity of results (**Figure 4D**) and trueness (**Figure 4E**).

Linearity, represented by R^2^ values, reflects the degree to which the observed overall labeled CHO abundances aligned with the expected values across the four spike levels (25 ng, 50 ng, 75 ng, and 100 ng). Despite some variation, the R^2^ values were generally high, suggesting that all tools achieved reasonably linear results. Byos displayed the highest R^2^ (0.962), followed by SpectroMine (0.910), Mascot (0.905), and FragPipe (0.904). PEAKS (0.896) and MaxQuant (0.888) exhibited slightly lower R^2^ values, indicating a minor reduction in linearity.

Trueness, assessed by absolute bias, revealed more pronounced differences in tool performance. Absolute bias was calculated as the deviation between the experimental overall labeled CHO protein abundance and the theoretical spike amount. Byos exhibited the highest trueness with the lowest mean absolute bias (8.9), followed by SpectroMine (14.5), Mascot (15.8), and PEAKS (19.5). FragPipe (20.3) and MaxQuant (21.3) showed higher absolute biases. Additionally, the range bars indicated that MaxQuant exhibited the greatest variability across spike levels, whereas Byos, SpectroMine, and Mascot demonstrated narrower ranges, with SpectroMine showing the most consistency.

Considering R^2^ values, mean absolute biases, and variability in bias, Byos outperformed the other tools, displaying high linearity and low mean bias across all spike levels. SpectroMine and Mascot followed closely with comparable metrics. This assessment was further visualized in a heatmap summarizing all evaluated metrics in this study (**Figure 4F**), where Byos and SpectroMine stood out for their strong overall performance. Additionally, PEAKS outperformed in protein coverage, while FragPipe exhibited superior data extraction consistency, achieving the highest count of quantifiable proteins and peptides.

## CONCLUSIONS

In this study, a rigorous benchmarking approach was employed to evaluate the qualitative and quantitative performance of six MS/MS search tools, specifically for monitoring HCPs in purified DS. By assessing these tools across varying spike levels, a detailed performance analysis was conducted, focusing on metrics such as peptide and protein identifications, triplicate consistency, and quantitative reliability after CV filtering and outlier removal. At the peptide level, statistical significance of log2FC values between spike levels was determined using unpaired Student’s t-tests, complemented by bootstrapping to assess differential linearity between observed and expected values. At the protein level, HMC modeling estimated posterior probability distributions, incorporating uncertainty into log2FC calculations. BDT was applied through utility maximization, generating scores that rewarded differential accuracy while penalizing high variability. Further evaluation focused on trueness and linearity, with particular attention to absolute biases and the alignment of observed values with theoretical expectations across spike levels. Tools achieving high utility scores, low bias, and good linearity emerged as top performers in terms of quantitative reliability. Ultimately, these findings underscore the distinct strengths of each MS/MS search tool: some excelled in peptide identifications and protein coverage, while others demonstrated superiority in critical quantitative metrics such as differential estimation, linearity, and trueness. This comprehensive benchmarking approach highlights the importance of tool selection based on specific research objectives. Whether the goal is extensive identification coverage, precise differential abundance analysis, or high-confidence HCP quantification, these findings provide valuable insights into the performance trade-offs inherent to each tool. For future LFQ applications in HCP monitoring, where precision and quantification fidelity are paramount, this study establishes a clear framework for selecting tools that meet these requirements. Unlike broader benchmarking studies that often assess proteins or peptides without stringent criteria, this approach applied multiple filters to distinguish between identified and quantifiable proteins, prioritizing measurement consistency and reliability over depth of coverage. This deliberate filtering strategy, though potentially sacrificing sensitivity, enhances quantification reproducibility, an essential consideration for the biopharmaceutical industry in monitoring critical quality attributes such as residual HCPs.

## ASSOCIATED CONTENT

### Supporting Information

Additional details and results.

## AUTHOR INFORMATION

### Funding Sources

The authors are full-time employees of the GSK group of companies with stock compensation.

### Author contribution

The authors contributed to the design of the study, acquisition and analysis of data, and manuscript development.

## Notes

### Competing Interest Statement

All authors are employees of the GSK group of companies.

### Summary of Updates

Figures 1, 2, and 5 were updated for clarity; a graphical abstract was added; supporting information file was revised; minor grammatical errors were corrected throughout the manuscript.

