## Supporting Information for "Comparative Analysis of MS/MS Search Algorithms in Label-Free Shotgun Proteomics for Monitoring Host-Cell Proteins Using Trapped Ion Mobility and ddaPASEF"

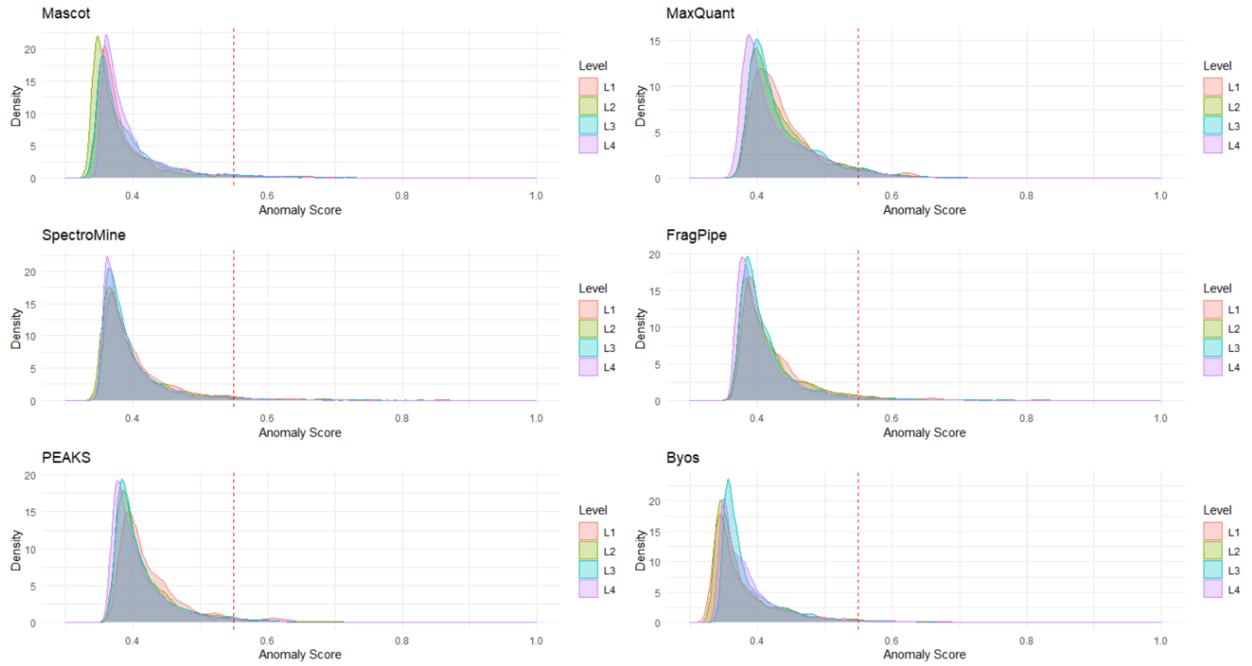


**Figure S1.** Density plots of the distribution of anomaly scores for peptide intensities analyzed with the Isolation Forest algorithm across six MS/MS search tools at four spike levels. An anomaly score threshold of 0.55 (indicated by the vertical red dashed line) was applied to identify outlier peptides, with scores exceeding this threshold classified as anomalies. Each spike level is represented by a different color, allowing for comparison of anomaly score distributions across concentration levels within each tool.

A

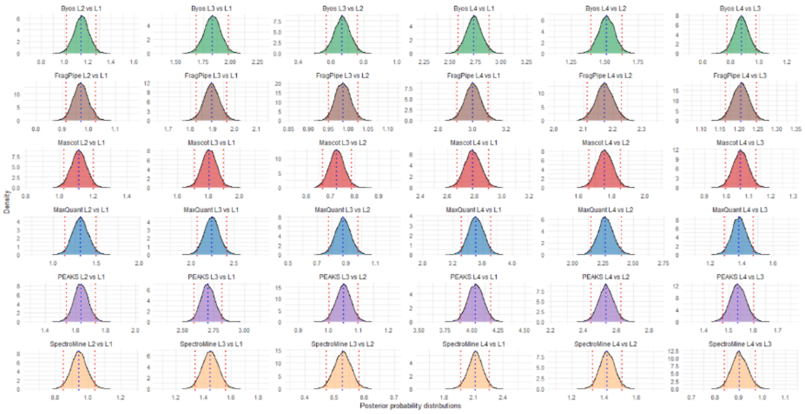

B

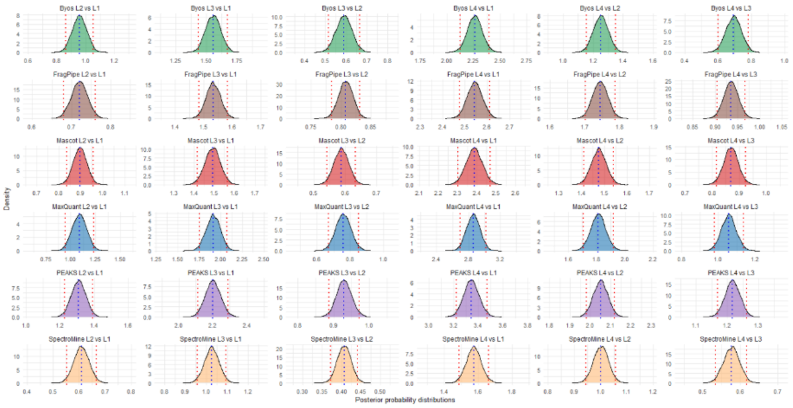

C

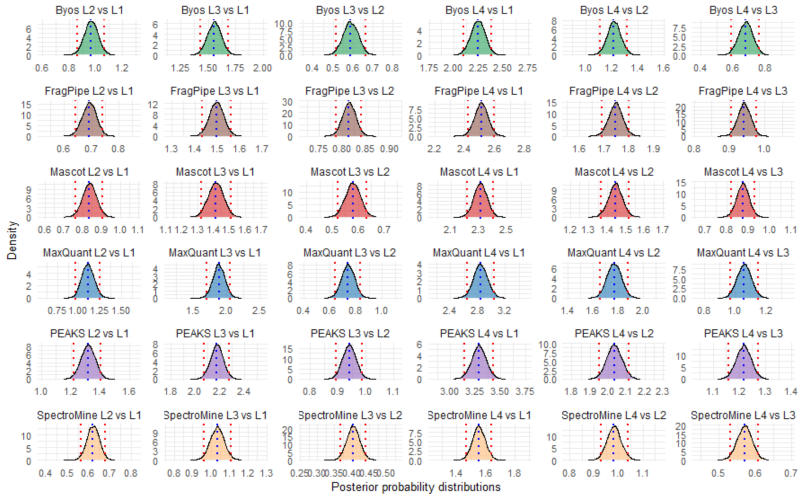


**Figure S2.** Posterior distributions of log2FC estimated by different MS/MS tools across various spike level comparisons using different protein aggregation methods, sum (A), average (B), and median (C). Vertical blue dashed lines indicate the posterior means. Vertical red dotted lines represent the 95% credible intervals, which provide the range within which the true FC is likely to lie with 95% probability. For SpectroMine, the sum aggregation demonstrated better agreement with the expected log2FC values, as reflected by narrower posterior distributions and lower σ, indicating this method's suitability for SpectroMine. In contrast, the other five tools performed better with the average or median aggregation methods.

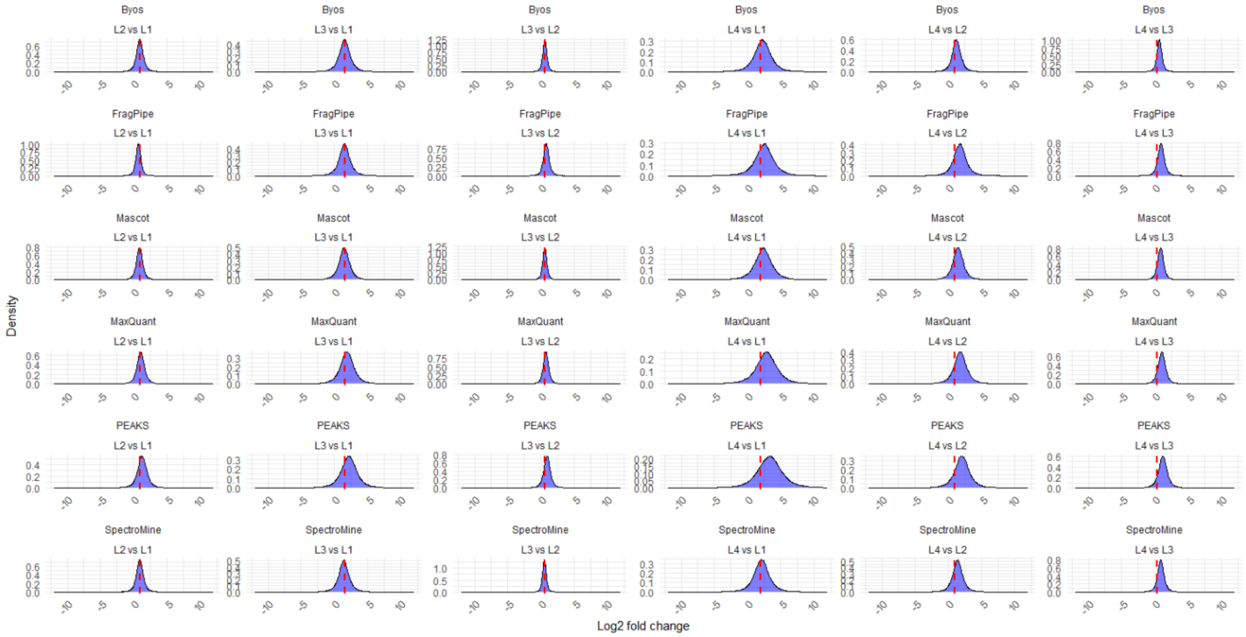

**Figure S3.** Density plots illustrate the posterior predictive distributions of log2FC for protein intensities across six MS/MS search tools over six pairwise comparisons (L2 vs. L1, L3 vs. L1, L3 vs. L2, L4 vs. L1, L4 vs. L2, and L4 vs. L3). Each plot shows the observed log2 fold change distribution (blue) overlaid with the posterior predictive distribution (red), providing a visual assessment of the fit between observed data and model predictions.

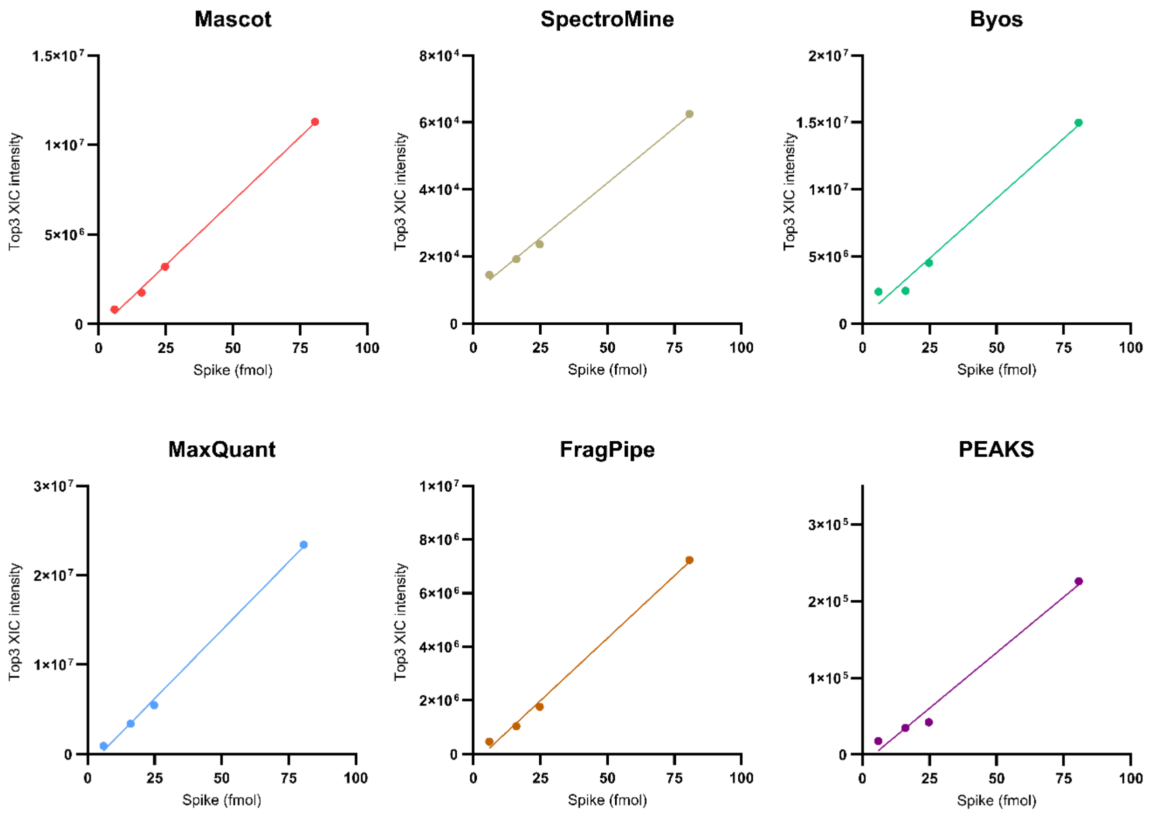


| | Mascot | MaxQuant | SpectroMine | FragPipe | PEAKS | Byos |
|---|---|---|---|---|---|---|
| R squared | 0.9981 | 0.9974 | 0.9957 | 0.9958 | 0.9829 | 0.9849 |

**Figure S4.** MassPREP regression plots showing the correlation between the spiked amount (fmol) of the four MassPREP digest standard proteins and their Hi3 generated intensities (using sum for SpectroMine, and average for the remaining tools).

**Table S1: Median CV% values of peptides identified by MS/MS tools as four spike levels**

| Spike level | Mascot | MaxQuant | SpectroMine | FragPipe | Byos | PEAKS |
|---|---|---|---|---|---|---|
| L1 | 16.6 | 34.8 | 14.5 | 11.7 | 14.4 | 18.1 |
| L2 | 16.8 | 32.1 | 13.6 | 10.9 | 16.2 | 15.3 |
| L3 | 19.5 | 29.6 | 14.2 | 12.6 | 18.3 | 17.9 |
| L4 | 18.4 | 23.4 | 14.5 | 11.4 | 18.7 | 15.7 |

**Table S2: Median absolute log2FC errors between expected and observed values of peptides for all six comparison levels**

| Spike level | Mascot | MaxQuant | SpectroMine | FragPipe | Byos | PEAKS |
|---|---|---|---|---|---|---|
| L2 vs. L1 | 0.20 | 0.21 | 0.39 | 0.26 | 0.21 | 0.47 |
| L3 vs. L1 | 0.23 | 0.35 | 0.57 | 0.27 | 0.33 | 0.80 |
| L3 vs. L2 | 0.13 | 0.18 | 0.17 | 0.30 | 0.20 | 0.37 |
| L4 vs. L1 | 0.63 | 1.07 | 0.49 | 0.80 | 0.59 | 1.63 |
| L4 vs. L2 | 0.63 | 0.96 | 0.27 | 1.01 | 0.55 | 1.30 |
| L4 vs. L3 | 0.55 | 0.73 | 0.25 | 0.68 | 0.40 | 0.91 |

**Table S3: Median log2FC errors between expected and posterior mean values of proteins for all six comparison levels**

| Spike level | Mascot | MaxQuant | SpectroMine | FragPipe | Byos | PEAKS |
|---|---|---|---|---|---|---|
| L2 vs. L1 | 0.16 | 0.12 | 0.08 | 0.47 | 0.06 | 0.39 |
| L3 vs. L1 | 0.09 | 0.27 | 0.13 | 0.05 | 0.03 | 0.47 |
| L3 vs. L2 | 0.25 | 0.51 | 0.08 | 0.35 | 0.18 | 0.74 |
| L4 vs. L1 | 0.01 | 0.38 | 0.15 | 0.47 | 0.01 | 0.67 |
| L4 vs. L2 | 0.56 | 0.86 | 0.50 | 0.80 | 0.32 | 1.04 |
| L4 vs. L3 | 1.08 | 1.37 | 1.14 | 1.19 | 0.76 | 1.57 |

**Table S4: Mean log10 posterior error probability values for all six comparison levels**

| Spike level | Mascot | MaxQuant | SpectroMine | FragPipe | Byos | PEAKS |
|---|---|---|---|---|---|---|
| L2 vs. L1 | 1.03 | 1.20 | 1.43 | 0.55 | 1.57 | 0.50 |
| L3 vs. L1 | 1.29 | 0.75 | 1.10 | 1.61 | 1.61 | 0.42 |
| L3 vs. L2 | 1.62 | 0.48 | 0.62 | 0.37 | 1.31 | 0.29 |
| L4 vs. L1 | 0.71 | 0.36 | 1.40 | 0.57 | 0.79 | 0.17 |
| L4 vs. L2 | 0.31 | 0.12 | 1.05 | 0.08 | 0.43 | -0.02 |
| L4 vs. L3 | -0.08 | -0.13 | 0.25 | -0.15 | 0.10 | -0.28 |
